# Carriers of human mitochondrial DNA macrohaplogroup M colonized India from southeastern Asia

**DOI:** 10.1101/047456

**Authors:** Patricia Marrero, Khaled K. Abu-Amero, Jose M Larruga, Vicente M Cabrera

**Affiliations:** School of Biological Sciences, University of East Anglia, Norwich NR4 7TJ, Norfolk, England; Glaucoma Research Chair, Department of Ophthalmology, College of Medicine, King Saud University; Riyadh, Saudi Arabia.; Departamento de Genetica, Universidad de La Laguna, La Laguna, Tenerife, Spain

**Author notes:** Corresponding author. E-mail address (V.M. Cabrera). Both authors equally contributed (Patricia Marrero, Khaled K. Abu-Amero). Actually retired (Vicente M Cabrera).

**Keywords:** Human evolution, mitochondrial DNA, out of Africa

## Abstract

**Objetives:** We suggest that the phylogeny and phylogeography of mtDNA macrohaplogroup M in Eurasia and Australasia is better explained supposing an out of Africa of modern humans following a northern route across the Levant than the most prevalent southern coastal route across Arabia and India proposed by others.

**Methods:** A total 206 Saudi samples belonging to macrohaplogroup M have been analyzed. In addition, 4107 published complete or nearly complete Eurasian and Australasian mtDNA genomes ascribed to the same macrohaplogroup have been included in a global phylogeographic analysis.

**Results:** Macrohaplogroup M has only historical implantation in West Eurasia including the Arabian Peninsula. Founder ages of M lineages in India are significantly younger than those in East Asia, Southeast Asia and Near Oceania. These results point to a colonization of the Indian subcontinent by modern humans carrying M lineages from the east instead the west side.

**Conclusions:** The existence of a northern route previously advanced by the phylogeography of mtDNA macrohaplogroup N is confirmed here by that of macrohaplogroup M. Taking this genetic evidence and those reported by other disciplines we have constructed a new and more conciliatory model to explain the history of modern humans out of Africa.

## Introduction

From a genetic perspective, built mainly on mtDNA data, the model of a recent African origin for modern humans (Cann et al., 1987; Vigilant et al., 1991) and their spread throughout Eurasia and Oceania replacing all archaic humans dwelling there, has held a dominant position in the scientific community. The recent paleogenetic discoveries of limited introgression in the genome of non-African modern humans, of genetic material from archaic, Neanderthal (Green et al., 2010; Prüfer et al., 2014) and Denisovan (Krause et al., 2010; Reich et al., 2011; Matthias Meyer et al., 2012) hominins has been solved adding a modest archaic assimilation note to the replacement statement (Stringer, 2014).

However, East Asia is a region where the alternative hypothesis of a continuous regional evolution of modern human from archaic populations is supported by the slow evolution of its Paleolithic archaeological record (Gao, 2013) and the irrefutable presence of early and fully modern humans in China, at least since 80 kya (Liu et al., 2010, 2015; Bae et al., 2014). Moreover, recently it has been detected ancient gene flow from early modern humans into Neanderthals from the Altai Mountains in Siberia at roughly 100 kya (Kuhlwilm et al., 2016). These data have negative repercussions on the phylogenetic hypothesis of a sole and fast dispersal of modern humans out of Africa around 60 kya, following a southern route (Macaulay et al., 2005; Thangaraj et al., 2005; Fernandes et al., 2012; Mellars et al., 2013). In principle, it could be adduced, as it was in the case of the early human remains from Skhul and Qafzeh in the Levant (Bar-Yosef, 1993), that the presence in China and Siberia of modern humans at that time was the result of a genetically unsuccessful exit from Africa. However, the fossil record in the area shows a clinal variation along a latitudinal gradient, with decreasing ages from China to Southeast Asia (Détroit et al., 2004; Barker et al., 2007; Mijares et al., 2010; Demeter et al., 2012) ending in Australia (Bowler et al., 2003). This is according to the mtDNA molecular clock but in an opposite way to the expected by the southern dispersal route. Clearly, the fossil record in East Asia would be more compatible with a model proposing an earlier exit from Africa of modern humans that arrived to China following a northern route, around 100 kya. Indeed, this northern route model was evidenced from the relative relationships obtained for worldwide human populations using classical genetic markers (Cavalli-Sforza et al., 1988; Nei and Roychoudhury, 1993) and archaeological record (Marks, 2005). Based on the phylogeography of mtDNA macrohaplogroup N, Maca-Meyer et al. (2001) inferred the existence of a northern route from the Levant that colonized Asia and carried modern humans to Australia for the first time. This idea was ignored or considered a simplistic interpretation (Palanichamy et al., 2004). On the contrary, the coastal southern route hypothesis has only received occasional criticism from the genetics field (Cordaux and Stoneking, 2003), and discrepancies with other disciplines are mainly limited to the age of exit from Africa of modern humans (Petraglia et al., 2012). However, subsequent research from the genetics, archaeology and paleoanthropology fields (Fregel et al., 2015), have given additional support to the possible existence of an early northern route followed by modern humans to colonize the Old World. At this respect, it is pertinent to mention that a recent whole-genome analysis comparing the ancient Eurasian component present in Egyptians and Ethiopians pointed to Egypt and Sinai as the more likely gateway in the exodus of modern humans out of Africa (Pagani et al., 2015). Furthermore, after a thoroughly revision of the evidence in support of a northern route signaled by mtDNA macrohaplogroup N (Fregel et al., 2015), we realized that the phylogeny and phylogeography of mtDNA macrohaplogroup M fit better in a northern route accompanying N than in a southern coastal route as was previously suggested (Maca-Meyer et al., 2001). In fact, M in the Arabian Peninsula, the source of our experimental data seems to have a recent historical implantation as in all western Eurasia. The unexpected detection of M lineages in Late Pleistocene European hunter-gatherers (Posth et al. 2016), possibly mirrors the back migration into Africa of haplogroup M1 that most probably arrived to Northern Africa through western Eurasia, in Paleolithic times (Olivieri et al., 2006; González et al., 2007; Pennarun et al., 2012). The founder age of M in India is younger than in eastern Asia and Near Oceania and so, southern Asia might better be perceived as a receiver more than an emissary of M lineages. In this study, we built a more conciliatory model for the history of Homo sapiens in Eurasia that might attract the reluctant East Asian position on the premises of only an early exit from Africa and only a sole northern route across the Levant, followed by early modern humans to colonize the Old World.

## Material and methods

### Sampling information

A total of 206 samples from unrelated Saudi healthy donors belonging to mtDNA macrohaplogroup M were analyzed in this study, 163 of them have been previously published (Abu-Amero et al., 2008); (Fregel et al., 2015). To fully characterize these M lineages, 17 complete mtDNA genomes from Saudi samples were sequenced. In addition, 7 unpublished complete mtDNA genomes, belonging to macrohaplogroup M, from preceding studies were included (Tanaka et al., 2004; González et al., 2007). Only individuals with known maternal ancestors for at least three generations were considered in this study. We included in the analysis 4107 published complete or nearly complete mtDNA genomes belonging to macrohaplogroup M of Eurasian and Oceania origin. To accurately establish the geographic ranges of the relatively rare M haplogroups, we screened 73,215 partial sequences from the literature as previously detailed in Fregel et al. (2015). Human population sampling procedures followed the guidelines outlined by the Ethics Committee for Human Research at the University of La Laguna and by the Institutional Review Board at the King Saud University. Written consent was recorded from all participants prior to taking part in the study.

### MtDNA sequencing

Total DNA was isolated from buccal or blood samples using the POREGENE DNA isolation kit from Gentra Systems (Minneapolis, USA).The mtDNA hypervariable regions I and II of the 43 new Saudi Arabian samples were amplified and sequenced for both complementary strands as detailed elsewhere (Bekada et al., 2013). When necessary for unequivocal assortment into specific M subclades, the 206 Saudi M samples were additionally analyzed for haplogroup diagnostic SNPs using partial sequencing of the mtDNA fragments including those SNPs, or typed by SNaPshot multiplex reactions (Quintáns et al., 2004). For mtDNA genome sequencing, amplification primers and PCR conditions were as previously published (Maca-Meyer et al., 2001). Successfully amplified products were sequenced for both complementary strands using the DYEnamic^TM^ETDye terminator kit (Amersham Biosciences) and samples run on MegaBACE^TM^ 1000 (Amersham Biosciences) according to the manufacturer's protocol. The 24 new complete mtDNA sequences have been deposited in GenBank under accession numbers KR074233-KR074256 (Figure S1).

### Previous published data compilation

Sequences belonging to specific M haplogroups were obtained from public databases such as NCBI, MITOMAP, the-1000 Genomes Project and from the literature. We searched for mtDNA lineages directly using diagnostic SNPs, or by submitting short fragments including those diagnostic SNPs to a BLAST search (http://blast.st-va.ncbi.nlm.nih.gov/Blast.cgi). Haplotypes extracted from the literature were transformed into sequences using the HaploSearch program (Fregel and Delgado, 2011). Sequences were manually aligned and compared to the rCRS (Andrews et al., 1999) with BioEdit Sequence Alignment program (Hall, 1999). Haplogroup assignment was performed by hand, screening for diagnostic positions or motifs at both hypervariable and coding regions whenever possible. Sequence alignment and haplogroup assignment was carried out twice by two independent researchers and any discrepancy resolved according to the PhyloTree database (van Oven and Kayser, 2009).

### Phylogenetic analysis

Phylogenetic trees were constructed by means of the Network v4.6.1.2 program using the Reduced Median and the Median Joining algorithms in sequent order (Bandelt et al., 1999). Resting reticulations were manually resolved. Haplogroup branches were named following the nomenclature proposed by the PhyloTree database (van Oven and Kayser, 2009). Coalescence ages were estimated by using statistics rho (Forster et al., 1996) and sigma (Saillard et al., 2000), and the calibration rate proposed by (Soares et al., 2009). Differences in coalescence ages were calculated by two-tailed t-tests. It was considered that the mean and standard error estimated from mean haplogroup ages calculated from different samples and methods were normally distributed.

### Global phylogeographic analysis

In this study, we are dealing with the earliest periods of the out-of-Africa spread. Given that subsequent demographic events most probably eroded those early movements, we omitted spatial geographic distributions of haplogroups according to their contemporary frequencies or diversities. The presence/absence of M basal lineages was used to establish the present-day haplogroup geographic range, while the overlapping geographic area of those haplogroups was considered as the most probable center of the old expansion. Pearson correlation tests were performed using geographic coordinates obtained by Google Earth software (https://earth.google.com).

### Geographic subdivision of India and regional haplogroup assignation

Attending only to geographic criteria, India was roughly subdivided into four different sampling areas: Northwest, including Kashmir, Himachal Pradesh, Punjab, Haryana, Uttarakhand, Rajasthan, Uttar Pradesh, Gujarat, and Madhya Pradesh states; Southwest, including Maharashtra, Karnataka and Kerala; Northeast including Bihar, Sikkim, Arunachal Pradesh, Assam, Nagaland, Meghalaya, Tripura, Jharkhand, West Bengal, Chhattisgarh and Orissa; and Southeast, represented by Andhra Pradesh, Tamil Nadu and Sri Lanka. We are very skeptical of the possibility that the actual genetic structure of India is the result of its original colonization, so the ethnic or linguistic affiliation of the samples were not considered but only its geographic origin. For the same motif we did not use the present day frequency and diversity of the haplogroups but their geographic ranges and radiation ages. The criteria followed to assign haplogroups to different regions within India were: consistent detection in an area (at least 90% of the cases in northern and 67% in southern areas) and absence or limited presence in the alternative areas (equal or less than 10% of the cases in northern and 33% in southern areas). We considered widespread those haplogroups consistently found in all the Indian areas and also found in surrounding areas as Pakistan or Iran at the west, Tibet or Nepal at the north, and Bangladesh or Myanmar at the east.

## Results and discussion

### Haplogroup M in western Eurasia with emphasis on Saudi Arabia

The lack of ancient and autochthonous mtDNA M lineages in western Eurasia is, at least, surprising if we think that the out of Africa colonization of the Old World by modern humans began through that region. Indeed, the presence of haplogroup M1 lineages in Mediterranean Europe and the Middle East has been explained as the result of secondary spreads of this haplogroup from northern Africa where it had a Paleolithic implantation (Olivieri et al., 2006; González et al., 2007; Pennarun et al., 2012). Likewise, the presence of eastern Asian M lineages belonging to C, D, G and Z haplogroups, mainly in Finno-Ugric-speaking populations of north and eastern Europe, seems to be the footprints of successive westward migration waves of Asiatic nomads occurred from Mesolithic period to historic times, as documented by the Magyar occupation of the Carpathian basin at the 9th century (Lahermo et al., 2000; Tambets et al., 2004; Ingman and Gyllensten, 2007; Nádasi et al., 2007; Tömöry et al., 2007; Lappalainen et al., 2008; Derenko et al., 2010, 2012; Der Sarkissian et al., 2013). South Asian influences on the west have been also detected by the presence of M4, M49 and M61 Indian lineages in Mesopotamian remains (Witas et al., 2013). In addition, ancestral mtDNA links between European Romani groups and northwest India populations were evidenced by the sharing of M5a1, M18, M25 and M35b lineages (Gresham et al., 2001; Kalaydjieva et al., 2001; Malyarchuk et al., 2008; Mendizabal et al., 2011; Gómez-Carballa et al., 2013).

Focusing on the mtDNA haplogroup M profile of Saudi Arabia, it represents about 7% of the Saudi maternal gene pool (Abu-Amero et al., 2007, 2008). Of the 206 M haplotypes sampled in Saudi Arabia (Table S1), 53% belong to the northern African M1 haplogroup, being both eastern African M1a and northern African M1b branches well represented (Table S1 and Figure S1). This fact contrasts with the sole presence of eastern M1a representatives in Yemen (Vyas et al., 2015). Therefore, most probably, M1b lineages reached Saudi Arabia from the Levant. M lineages with indubitable Indian origin account for 39% of the Saudi M pool, whereas the resting 8% has, most probably, a southeastern Asian source. The geographic origin of the Indian contribution seems not to be biased as 53% of the lineages might be assigned to eastern and 47% to western Indian regions (Metspalu et al., 2004; Chandrasekar et al., 2009; Maji et al., 2009). However, the fact that the two M lineages characteristic of the Andaman aborigines (Endicott et al., 2003; Thangaraj et al., 2005; Barik et al., 2008) are present in the Saudi sample deserves mention. The isolate Ar2461 (Table S1) has the diagnostic mutations of the Andaman branch M31a1 in the regulatory (249d, 16311) and in the coding (3975, 3999) regions. On the other hand, the complete mtDNA sequence of Ar1076 (Figure S1) belongs to the M32c Andamese branch, matching another complete sequence reported for Madagascar (Dubut et al., 2009). It must be stressed that this branch has been steadily found in all mtDNA reports on Madagascar (Hurles et al., 2005; Tofanelli et al., 2009; Capredon et al., 2013). Although at first these haplogroups were taken as evidence that the indigenous Andamanese represent the descendants of the first out-of-Africa dispersal of modern humans (Endicott et al., 2003; Thangaraj et al., 2005), more recent studies support a late Paleolithic colonization of the Andaman Islands (Palanichamy et al., 2006; Wang et al., 2011). About the possible origin of these settlers, it must be mentioned that different branches of M31 have been found in northeastern India, Nepal and Myanmar (Thangaraj et al., 2006; Reddy et al., 2007; Barik et al., 2008; Fornarino et al., 2009; Wang et al., 2011). Inasmuch as M32c, it has been also detected in Indonesia (Hill et al., 2007) and in Malaysia (Haslindawaty et al., 2010; Maruyama et al., 2010). Another interesting link involving India, Saudi Arabia and the Mauritius Island is the case of haplogroup M81,defined in PhyloTree.org Build16 (van Oven and Kayser, 2009). It was first detected as a sole sequence in a LHON patient from India (Khan et al., 2013). The Saudi sample Ar567 shares with this Indian sequence substitutions 215, 4254, 6620, 13590, 16129 and 16311 and, in addition, it shares substitutions 151, 6170, 7954 and 16263 with a complete sequence from the Mauritius Island (Fregel et al., 2014). At first sight, four different mutations occurring in a short segment of 11 bp, from positions 5742 to 5752, differentiate the Mauritius from the Saudi sequence. However, a closer inspection reveals they represent only two additional shared mutations differently interpreted: the loss by transversion to G of a C at position 5743 in the Ma12 reading corresponds to a deletion of C in the same position in the Ar567 lecture, and the loss of a G by transition to A at position 5746 in Ma12 corresponds to an A insertion at position 5752 in Ar567. Therefore, both sequences can be only distinguished by transition 12522 present in the Saudi sample and absent in Ma12 (Figure S1). Curiously, these affinities between samples from Saudi Arabia and those from Indian Ocean islands near the African shores can be extended to the Saudi Q1a1 sequence Ar196 (Table S1), which has exact matches only with MA405 sample from Madagascar (Capredon et al., 2013), and to the presence in Saudi Arabia (Fregel et al., 2015) and Mauritius (Fregel et al., 2014) of different lineages belonging to the Indian branch (M42b) for which a deep link with the Australian M42a branch has been detected (Kumar et al., 2009). Other lineages present in the Saudi mtDNA pool, as M20, E1a1a1 or M7c1 point to specific arrivals from southeastern Asia. Interesting as these affinities and provenances are, the fact that all these lineages represent isolates in Saudi Arabia having matches or being very related to those found in the original areas strongly support the suggestion that they have been incorporated to the Saudi pool in historic times as result of the Islamic expansion and the import of indentured workers from India and southeastern Asia by the European colonizers in the past and by the own Saudi at present times (Abu-Amero et al., 2008). In conclusion, Saudi Arabia lacks of ancient and autochthonous M haplogroups likewise the rest of western Eurasia.

### About the origin of the North African haplogroup M1

The existence of haplogroup M lineages in Africa was first detected in Ethiopian populations by RFLP analysis (Passarino et al., 1998). Although an Asian influence was contemplated to explain the presence of this M component on the maternal Ethiopian pool, the dearth of M lineages in the Levant and its abundance in south Asia gave strength to the hypothesis that haplogroup M1 in Ethiopia was a genetic indicator of the southern route out of Africa. In addition, it was pointed out that probably this was the only successful early dispersal (Quintana-Murci et al., 1999). However, the limited geographic range and genetic diversity of M in Africa compared to India was used as an argument against this hypothesis (Maca-Meyer et al., 2001; Roychoudhury et al., 2001; Metspalu et al., 2004; Olivieri et al., 2006; Thangaraj et al., 2006; González et al., 2007), instead proposing M1 as a signal of backflow to Africa, most probably from the Indian subcontinent. However, after extensive phylogenetic and phylogeographic analyses for this marker (Metspalu et al., 2004; Olivieri et al., 2006; Sun et al., 2006; González et al., 2007; Pennarun et al., 2012), this supposed India to Africa connection was not found. The detection in southeast Asia of new lineages that share with M1 the 14110 substitution (Kong et al., 2011; Peng et al., 2011), gave rise to the definition of a new macrohaplogroup named M1'20'51 by PhyloTree.org Build 16 (van Oven and Kayser, 2009). However, this substitution is not an invariable position (Table S2) and, therefore, its sharing by common ancestry is not warranted (Pennarun et al., 2012). Besides, we realized that, in addition, haplogroups M1 and M20 share transition 16129 which, although highly variable, would add support to this basal unification as the most parsimonious phylogenetic reconstruction (Figure S1). However, after that, a recent study reported a new mtDNA haplogroup from Myanmar, named M84, that also roots at the basal node represented by transition 14110 (Li et al., 2015). This haplogroup shares with M20 the transition 16272 which is more conservative than 16129 (Soares et al., 2009), therefore, weakening any specific relationship between haplogroups M1 and M20 beyond its common basal node. With the exception of M84, that seems to be limited to Myanmar, India and Southern China populations (Li et al., 2015), the phylogeography of these haplogroups is extend and complex (Table S3). M1 is found from Portugal and Senegal in the west to the Caucasus, Pakistan and Tibet at the east and, from Guinea-Bissau and Tanzania in the south to Russia at the north (Kivisild et al., 2004; Olivieri et al., 2006; Gonder et al., 2007; González et al., 2007, 2007; Zhao et al., 2009; Malyarchuk et al., 2010; Pennarun et al., 2012; Yunusbayev et al., 2012; Siddiqi et al., 2014) but, its highest diversity is found in Ethiopia and the Maghreb as the isolates detected at the borders are lineages derived from M1a branch in Russia and Tanzania and M1b branch in Guinea Bissau and Tibet. The geographic range of M20 and M51 largely overlaps, showing a common wide area in southeastern Asia including Myanmar and Malaysia at the west, Philippines and Hainan at the east and Tibet at the northwest (Table S3). In addition, for M20 the western border further stretches to Bangladesh (Gazi et al., 2013) and even to Assam in India (Cordaux et al., 2003) and the eastern border of M51 to Fujian and Taiwan (Kong et al., 2011). The primary split, separating the ancestors of these four haplogroups occurred around 56 kya, coinciding with a warm climate period so that this split could have occurred in any place of their common geographic ranges although, most probably, in its core area. Thus, it might be possible that the birth of the M1 ancestor was in southeastern Asia instead of India. Based on the scarcity and low diversity of M1 along the southern route (Vyas et al., 2015), and archaeological affinities between Levant and North Africa (Olivieri et al., 2006; González et al., 2007), it has been suggested that the route followed by the M1 bearers to reach Africa was across the Levant not to the Bab el Mandeb strait. However, the actual representatives of M1, from the Levant to the Tibet, are derived lineages with ancestors in Africa. Until now, basal lineages of M1 have not been detected in any of the northern or southern hypothetical paths. The only indirect support for a Levantine route of M1 is the fact that all the other Eurasian lineages as N1, X1 or U6, that returned to Africa in secondary backflows, had their most probable origins in central or western Eurasia instead of India.

### The role of India in the origin and expansion of macrohaplogroup M

Macrohaplogroup M in South Asia is characterized by a great diversity, a deep coalescence age, and the autochthonous nature of its lineages. These characteristics have been used in support of an in-situ origin of the M Indian lineages and a rapid dispersal eastwards to colonize southeastern Asia following a southern coastal route (Macaulay et al., 2005; Thangaraj et al., 2005; Chandrasekar et al., 2009). However, there are several arguments against this hypothesis. First, as geographic barriers seem not to be stronger at the western than at the eastern border of the subcontinent (Boivin et al., 2013), it should be expected that macrohaplogroup M would simultaneously radiated to western as well as to eastern areas if India was its first center of expansion. However, while at the east its expansion crossed the Ganges and quickly reached Australia, at the west it seems to have found an insurmountable barrier at the Indus bank as M frequencies suddenly drop from 65.5 ± 4.3% in India to 5.3 ± 1.0 % in Iran (p < 0.0001) (Metspalu et al., 2004; Ashrafian-Bonab et al., 2007; Farjadian et al., 2011; Derenko et al., 2013). Furthermore, unlike southeastern Asia and Australia, autochthonous M lineages have not been detected to the west of South Asia. It might be adduced that later expansions from the West replaced the M lineages in those regions, however, massive expansions of western mtDNA lineages in India have not been detected, instead it has been pointed out that most of the extant mtDNA boundaries in South and Southwest Asia were likely shaped during the initial settlement of Eurasia by anatomically modern humans (Metspalu et al. 2004); only a small fraction of the Caucasoid specific mtDNA lineages found in Indian populations can be ascribed to a relatively recent admixture (Kivisild et al. 1999). Secondly, whenever it has been compared, the founder age of M in India is significantly (p = 0.002) younger than those in Eastern and Southeastern Asia and Australo-melanesian centers (Table 1). It has been argued that this could be due to an uneven distribution and sampling of M lineages in India (Sun et al., 2006; Thangaraj et al., 2006). However, when the same weight is given to every lineage, independently of their respective abundance (Table 1), the founder age slightly diminish in India and raises in East Asia compared to their respective weighted founder ages. Recently, it has been admitted that the initial radiation of macrohaplogroup M could have occurred in eastern India or further east following the southern route (Mellars et al., 2013). In fact, the possibility that the first radiation of M were in East Asia instead of India had been proposed by Maca-Meyer et al. (2001). We think that it would be a hard task to detect today the original focus of macrohaplogroup M expansion into India if it really occurred. Indeed, we noticed that the mean radiation age (30.1 ± 9.3 ky) of the M haplogroups that could be considered of widespread implantation in India is significantly older (Two-tailed p value = 0.0147) than the mean radiation age (22.2 ± 9.6 ky) of those with more localized ranges (Table S3). However, comparisons between mean radiation ages of Indian M haplogroups with eastern vs western (p = 0.275) and northern vs southern (p = 0.580) geographic ranges are not statistically significant. Furthermore, it must be taken into account the possibility that some M haplogroups with dominant implantation in northeastern India, as the case of M49, are in fact secondary radiations from Myanmar (Summerer et al., 2014; Li et al., 2015). It is evident that the founder age of macrohaplogroup M increases eastwards from South Asia. Not only is the founder age of M in East Asia significantly greater than in South Asia but, the former is younger than the one estimated for southeast Asia although the probability value (p = 0.0998) does not reach significance probably because the estimation for the area calculated by Sun et al. (2006) was previous to the detection of M clades with very deep ages in southeast Asia (Kong et al., 2011; Peng et al., 2011; Zhang et al., 2013). In turn, the founder age for M in Oceania is significantly older than the ones estimated for East Asia (p = 0.004) and for Southeast Asia (p = 0.032). This decreasing age gradient moving westwards is in accordance with the hypothesis that carriers of macrohaplogroup M lineages colonized India from the East instead the West. It could be argued that South Asia was colonized very early by the southern eastward coastal expansion of modern humans out of Africa but those primitive mtDNA lineages were extinguished by genetic drift and the subcontinent recolonized latter from eastern groups left on the way to Australia. However, this argument is in contradiction, first, with the suggestion, based on past population size prediction deduced from mtDNA variation, that approximately 45 and 20 kya most of humanity lived in southern Asia (Atkinson et al., 2008), and second, with the mean founder age of macrohaplogroup R in South Asia (62.5 ±3.5 ky) that is significantly older (p = 0.004) than its M counterpart (45.7 ± 7.8 ky). These data are in agreement with a colonization of South Asia by two independent waves of settlers. Based on the phylogeny and phylogeography of haplogroups M and U in India, this two-wave model was already proposed in the first mtDNA studies of Indian populations (Kivisild et al., 1999). Curiously, this idea was abandoned in favor of a single southern route simultaneously carrying the three Eurasian mtDNA lineages M, N and R. Now, if the born of macrohaplogroup M at some place between southeast Asia and near Oceania is accepted, the putative connection between Australia and India, based on the nearly simultaneous radiation of branches M42a and M42b in both areas respectively (Kumar et al., 2009), could be explained as the result of an expansion from a nearly equidistant center of radiation, not as a directional colonization from India to Australia following a southern route. The tentative junction of M42 with the specific Southeast Asian M74 lineage, based on a shared transition at 8251 in PhyloTree.org Build 16 (van Oven and Kayser, 2009) gives additional support to this hypothesis.

**Table 1.**
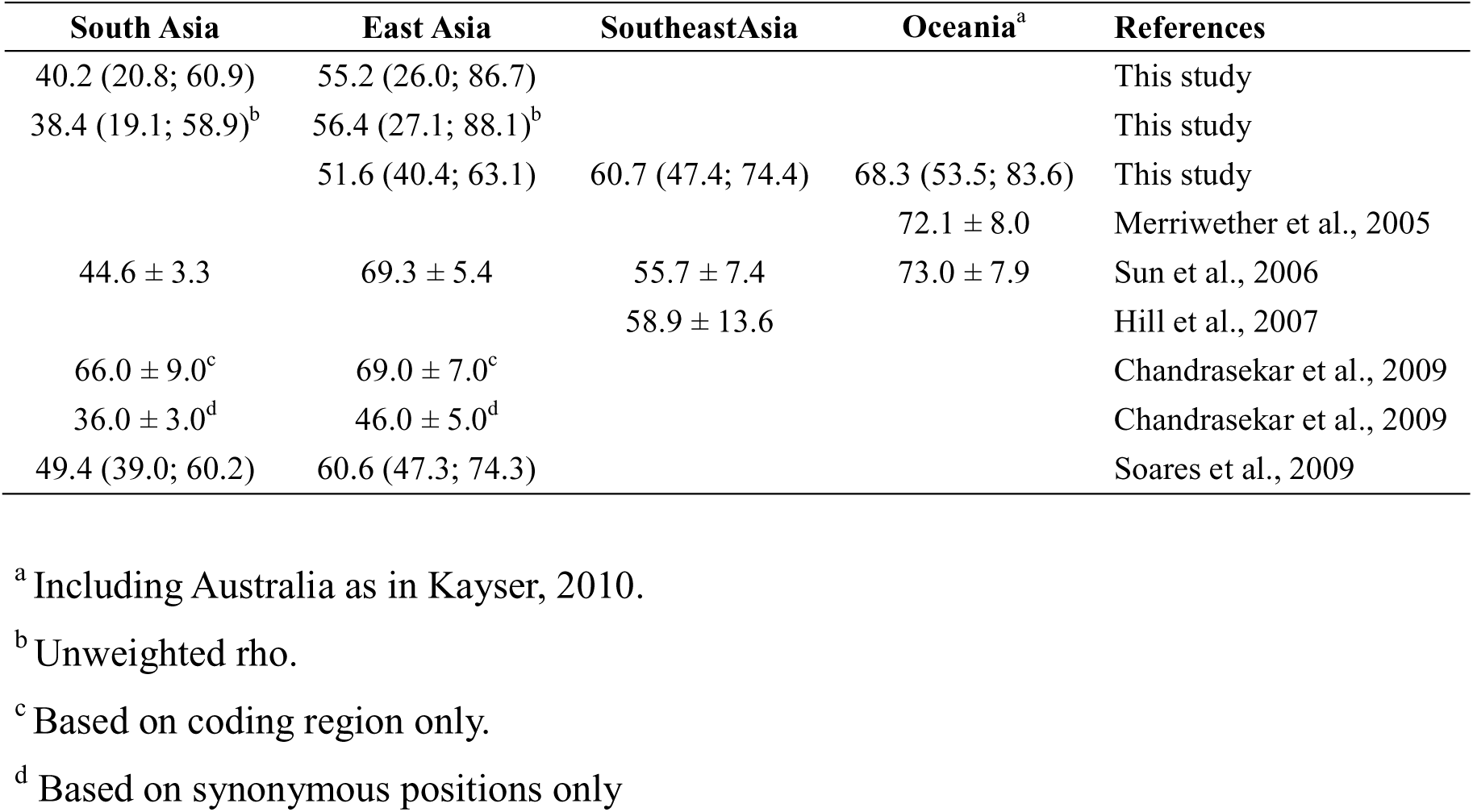
Founder ages (kya) for macrohaplogroup M in South and East Asia.

| South Asia | East Asia | SoutheastAsia | Oceania<sup>a</sup> | References |
| --- | --- | --- | --- | --- |
| 40.2 (20.8; 60.9) | 55.2 (26.0; 86.7) |  |  | This study |
| 38.4 (19.1; 58.9)<sup>b</sup> | 56.4 (27.1; 88.1)<sup>b</sup> |  |  | This study |
|  | 51.6 (40.4; 63.1) | 60.7 (47.4; 74.4) | 68.3 (53.5; 83.6) | This study |
|  |  |  | 72.1 ± 8.0 | Merriwether et al., 2005 |
| 44.6 ± 3.3 | 69.3 ± 5.4 | 55.7 ± 7.4 | 73.0 ± 7.9 | Sun et al., 2006 |
|  |  | 58.9 ± 13.6 |  | Hill et al., 2007 |
| 66.0 ± 9.0<sup>c</sup> | 69.0 ± 7.0<sup>c</sup> |  |  | Chandrasekar et al., 2009 |
| 36.0 ± 3.0<sup>d</sup> | 46.0 ± 5.0<sup>d</sup> |  |  | Chandrasekar et al., 2009 |
| 49.4 (39.0; 60.2) | 60.6 (47.3; 74.3) |  |  | Soares et al., 2009 |

<sup>a</sup> Including Australia as in Kayser, 2010.
<sup>b</sup> Unweighted rho.
<sup>c</sup> Based on coding region only.
<sup>d</sup> Based on synonymous positions only

### Basal M superhaplogroups involving South Asia

As the most parsimonious topology, the existence of several M superhaplogroups, embracing different lineages based on one or a few moderately recurrent mutations have been recognized in PhyloTree.org Build 16 (van Oven and Kayser, 2009). Following this trend we have, provisionally, unified haplogroup M11 and the recently defined haplogroup M82 (Li et al., 2015) under macrohaplogroup M11'82 because both share the very conservative transition 8108 (Table S2). Attending to their phylogeography these superhaplogroups usually link very wide geographic areas (Table S2). In accordance with our suggestion that the carriers of Macrohaplogroup M colonized South Asia later than southeastern Asia and Oceania and that this mtDNA gene flow had an eastern origin we have observed that the mean radiation age of those superhaplogroups involving South Asia (39.70 ± 3.24 ky) is significantly younger (p = 0.003) than the one relating East and Southeast Asia (55.60 ± 2.94 ky). In this comparison we considered the South Asia-Southeast Asia-Australian link deduced from superhaplogroup M42'74 as specifically involving India. However, we considered superhaplogroup M62'68 as indicative of an East Asia-Southeast Asia link because M62 has been found consistently in Tibet (Zhao et al., 2009; Qin et al., 2010; Qi et al., 2013) but only sporadically in northeast India (Chandrasekar et al., 2009).

### The perspective from other genetic markers

Based on the absence of autochthonous mtDNA haplogroup N lineages in India and the very deep age of N lineages in southeastern Asia we have, recently, reasserted (Fregel et al., 2015) our long ago formulated hypothesis that mtDNA macrohaplogroup N reached Australia following a northern route (Maca-Meyer et al., 2001). This time, we have found additional support from Paleogenetics that has demonstrated introgression in the genome of modern humans of DNA from Neanderthals (Green et al., 2010; Prüfer et al., 2014) and Denisovans (Reich et al., 2010; M. Meyer et al., 2012) hominins that, most probably, had northern geographic ranges (Sawyer et al., 2015). For the specific case of India, we proposed that the N and R lineages present in the subcontinent arrived, as secondary waves, from the north following the Indus and Ganges-Brahmaputra banks. Thus, in our opinion, India was first colonized by modern humans from two external geographic centers situated at the northwest and northeast borders of this subcontinent (Fregel et al., 2015). The same picture was depicted, long ago, by other authors also based on mtDNA variation in India (Kivisild et al., 1999), although suggesting an eastward southern route arrival for macrohaplogroup M, the same way proposed by Maca-Meyer et al. (2001). However, due to the lack of any mtDNA genetic evidence in support of an early migration along the southern route, we now propose an early exit of modern humans from Africa by the Levant and a unique northern route up to the Altai Mountains and, obliged by harsh weather, down to southern China and beyond (Fregel et al., 2015). Strong evidence, supporting the primitive colonization of India by modern humans in two waves, has come from genome wide analysis. After analyzing 132 samples, from 15 Indian states, for more than 500,000 autosomal SNPs, Reich et al. (2009) detected the existence of two ancient populations, genetically divergent, in India. One, the ancestral northern Indians, close to Middle Eastern, Central Asians and Europeans, and the other, the ancestral southern Indians, that is as distinct from the northern component and East Asians as they are from each other. Actually, the southern component is most prevalent in the Andamanese. Of particular interest is the fact that when populations of Southeast Asia and Near Oceania were incorporated to these genome wide analyses, the Andaman Islanders showed a closer affinity with southeast rather than South Asian populations (Chaubey and Endicott, 2013; Aghakhanian et al., 2015). It is evident that our mtDNA interpretation and the autosomal results give a very similar picture. Furthermore, independent support for the existence of an early center of primitive modern humans in southeastern Asia, that originated very early expansions, has come recently from improved phylogenetic resolution analysis of the Y-chromosome K-M526 haplogroup. It has been detected a rapid diversification process of this haplogroup in Southeast Asia-Oceania with subsequent westward expansions of the ancestors of haplogroups R and Q that make up the majority of paternal lineages in Central Asia and Europe (Karafet et al., 2014).

### The fossil evidence

There is strong contradiction between the paleontological and the genetic interpretations about the origin of modern human in East Asia. New and reliable chronometric dating techniques applied to morphologically classify human remains have demonstrated the presence of early and fully modern humans in southern China at least since 80 kya which is in support of regional continuity, or in situ evolution, of modern humans in East Asia (Smith and Spencer, 1984; Liu et al., 2015). On the other hand, no apparent genetic contribution from earlier hominids was detected in the maternal (Tanaka et al., 2004; Kong et al., 2011) and the paternal (Ke et al., 2001; Shi et al., 2005; Wang and Li, 2013) genetic pools of extant East Asian populations which was taken as in support of a recent replacement of archaic humans by modern African incomers in East Asia. Although recently, genome analysis have detected introgression of Neanderthal (Prüfer et al., 2014) and Denisovan (Matthias Meyer et al., 2012) DNA to the extant (Skoglund and Jakobsson, 2011) and the ancient (Fu et al., 2013a) genomes of modern humans in East Asia, this genetic contribution can be explained as a limited assimilation episode. Ancient DNA analysis of a morphologically early modern human from Tianyuan cave, in northeast China (Fu et al., 2013a) and that of a 45 ky old modern human from western Siberia (Fu et al., 2014) evidenced that both were bearers of B and U mtDNA lineages that are branches of haplogroup R which, in its turn, derives from macrohaplogroup N indicating that around 45 kya, people in North Asia carried already fully modern mtDNA lineages which is a hint of a northward secondary expansion of modern humans at that time. On the other hand, dates of the East Asian fossil record (Table S4) show a significant (p = 0.0008) positive correlation (r = 0.772) with their respective latitudes southward from China around 100 kya. This is in agreement with our proposition that the out of Africa of modern humans occurred across the Levant and that the Skhul and Qafzeh fossil remains in Israel were the first landmarks of that successful exit (Fregel et al., 2015). Subsequent evolution of those early modern humans in Asia might reconcile the replacement and continuity models into an inclusive synthesis. On the mtDNA side, the main problem for this reconciliation would be an strict adhesion to the chronological upper bound marked for the African exit by the coalescent age of macrohaplogroup L3 (Mellars et al., 2013). However, we think that mtDNA dating methods are still not reliable in absolute terms, mainly because we need accurate independent calibration for deep nodes and, in addition to selection, to take into account the effects of demographic parameters on the temporal variation of the substitution rate. Turning to the probable existence of a primitive center of modern human expansion in southeast Asia, as proposed here on the basis of mtDNA haplogroup M phylogeography, the existence of a positive westward longitudinal gradient (r = 0.551) of the fossil record dates in southeastern Asia, with the older ages in Philippines and the youngest in Sri-Lanka (table S4), deserves mention although, this time, the correlation is not statistically significant (p = 0.199), most probably due to the lack of Paleolithic fossil evidence in Myanmar and India. Certainly, this counter-clock trend has been perceived by other authors as an argument against the straight forward recent out of Africa dispersal of *Homo sapiens* model (Boivin et al., 2013).

### The archaeological evidence

The archaeological record in East Asia also seems to be in support of the regional continuity model. The persistence and dominance of simple core-flake assemblages throughout the Paleolithic in this area sharply contrasts with the Upper Paleolithic technical and cultural innovations in western Eurasia (Bar-Yosef, 2002). Different lithic assemblages with potential resemblances in Africa and in southeastern Asia are crucial to interpret the southern dispersal route of modern humans across India. Blade-microblade and core-flake technologies are both present in the Indian subcontinent. They are usually contemporary and, in some cases, core-flake industries as the Soanian even post-date the former (Mishra, 2008). Microlithics in India have been dated around 35-30 kya in southern India and in Sri Lanka (Dennell and Petraglia, 2012) but, recently, it has been established that the microblade technology presents continuity in central India since 45 kya (Mishra et al., 2013). Some authors have proposed that the Indian microlithics had an African original source (Mellars et al., 2013). This model has problems to explain the absence of microblades around that time eastwards the subcontinent or the chronological gap between the oldest microlithic dates in India and the arrival of modern humans to Australia. Other authors consider microlithics in India as local innovations (Dennell and Petraglia, 2012). However, the absence of an earlier blade technology in the Indian Paleolithic record makes an indigenous development unlikely (Mishra et al., 2013). Finally, from the age of the microblade technology in India, other authors deduced that modern humans skirted the Indian subcontinent in the first dispersal out of Africa taken a northerly route through the Middle East, Central Asia, and southeastern Asia across southern China (Mishra et al., 2013). For these authors, modern humans did not actually enter India until the time marked by the glacial climate of MIS 4. On the other hand, some core and flake industries in India have been considered as a link between those present in sub-Saharan Africa, Southeast Asia and Australia (Petraglia et al., 2007; Clarkson et al., 2009; Haslam et al., 2010). Core sites with ages around 77 ky in India would be compatible with an early dispersal of modern humans from Africa. This model confront problems as a later mtDNA molecular clock boundary proposed by some geneticists (Fernandes et al., 2012) and the lack of contemporary fossil record to confirm that this primitive technology was manufactured by modern humans. Finally, some Indian Late Pleistocene core-flake types as the Soanian seem to be related to similar industries as the Hoabinhian in southeastern Asia (Gaillard et al., 2010). We suggest that microlithic technology could signal the arrival of mtDNA macrohaplogroup R to India, following the northwest passage, as a branch of a global secondary dispersal of modern humans from some core area in west/central Asia than also affected Europe (Marks, 2005) and later reached North China across Siberia and Mongolia (Qu et al., 2013). On the other hand, macrohaplogroup M came to India from Southeast Asia following the Northeast Passage and carrying with them a simple core-flake technology.

## Conclusions

A new and integrative model, explaining the time and routes followed by modern humans in their exit from Africa is proposed in this study (Figure 1). First, we think that the exit from Africa followed a northern route across the Levant and that the fossils of early modern humans at Skhul and Qafzeh could be signals of this successful dispersal. These first modern humans carried mtDNA L3 undifferentiated lineages and brought primitive core-flake technology to Eurasia (Marks, 2005). The dates estimated for Skhul and Qafzeh remains (Table S4) are slightly out of the range calculated for the age of mtDNA macrohaplogroup L3 (78.3, 95% CI: 62.4;94.9 kya) based on ancient mtDNA genomes (Fu et al., 2013b). However, they are in accordance with the presence of modern and early modern humans in China around 100 kya, and with their subsequent presence in southeastern Asia about 70 kya. If we add the fact, also based on ancient genomes, that around 45 kya mtDNA lineages in northern Asia already belonged to derived haplogroups B and U (Fu et al., 2013a, 2014), we opine that are the geneticists who should synchronize its mtDNA molecular clock with the fossil record and not the reverse. Second, those early modern humans went further northwards, some at least to the Altai Mountains, and in the way they occasionally mixed with other hominids as Neanderthal and Denisovans. Harsh climatic conditions dispersed them southwards erasing the mtDNA genetic footprints of this pioneer northern phase (Fregel et al., 2015). Third, the surviving small groups already carried basic N and M mtDNA lineages. One of them, with only N maternal lineages, spread southwards to present-day southern China and probably, across the Sunda shelf reached Australia and the Philippines (Fregel et al., 2015). Fourth, other dispersed groups were the bearers of other N branches, including macrohaplogroup R, that did not reach Australia in the first wave, but enter India from the north, carrying with them the blade-microblade technology detected in this subcontinent, that also spread with other N and R branches to northern and western Eurasia, reaching Europe, the Levant and even northern Africa. Fifth, short after, another southeastern group, carrying undifferentiated M lineages, radiated from a core area, most probably localized in southeast Asia (including southern China), reaching India westwards and near Oceania eastwards. These haplogroup M bearers brought with them at least one of the primitive core-flake technologies present in India that, therefore, had to have a southeastern Asian origin. Sixth, in subsequent mild climatic windows, demographic growth dispersed macrohaplogroup M and N northwards, most probably from overlapping areas that in time colonized northern Asia and the New World.

**FIGURE 1.**
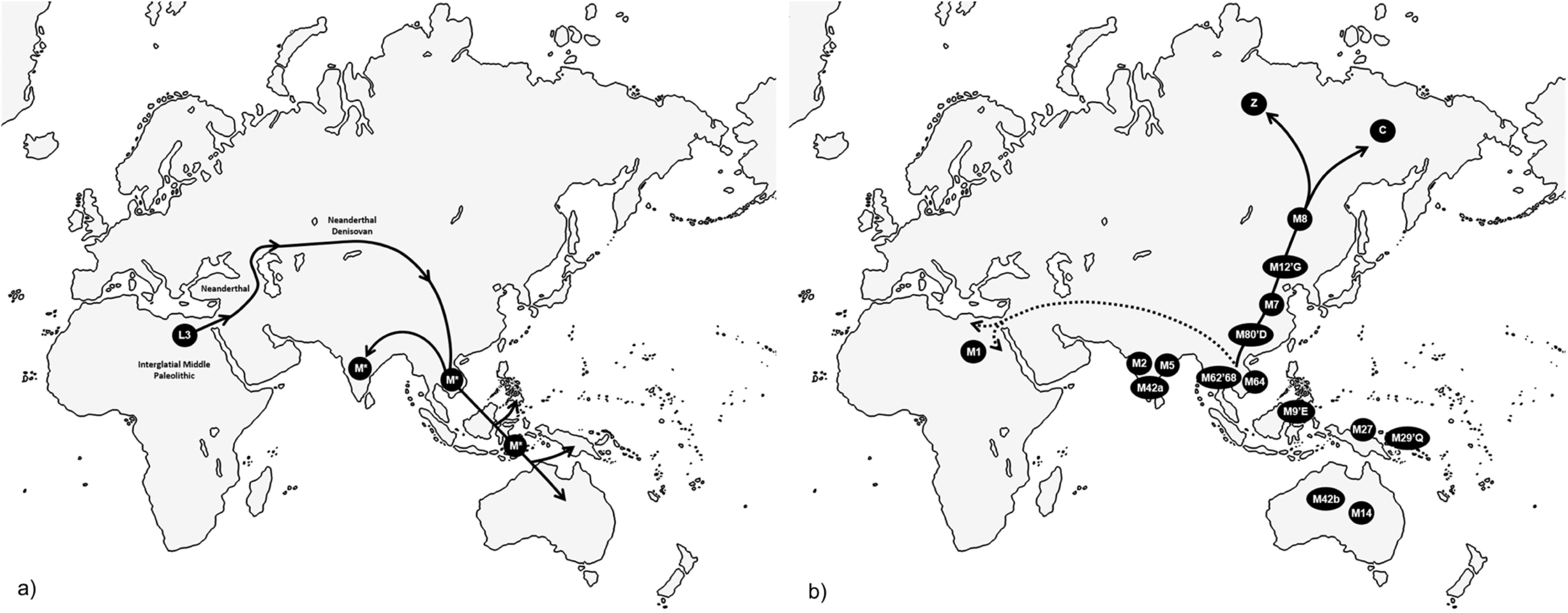
Proposed routes followed by modern humans in their exit from Africa: (a) Northern route to reach South Asia, the Philippines and nearly Oceania, and (b) secondary expansions northward through Asia to the Americas and southwest to North Africa and Europe.

It seems that under archaeological grounds there are data in support of a southern route exit from Africa and across the Arabian Peninsula and the Indian subcontinent (Petraglia et al., 2007, 2012; Clarkson et al., 2009; Rose, 2010; Armitage et al., 2011; Dennell and Petraglia, 2012; Scerri et al., 2014; Groucutt et al., 2015), but the lack of coetaneous fossil record leaves the hominid association to these stone tools unresolved. Even if they were modern humans they did not leave any trace in the mtDNA gene pool of the extant populations of Arabia or India. However, there is no necessity to invoke the existence of a southern route to interpret the landscape depicted by the mtDNA phylogeny and phylogeography.

## Acknowledgements

We are grateful to Dr. Ana M. González for her contribution and helpful comments to this work. P. Marrero was supported by a postdoctoral grant from the Spanish Education Ministry (EX2009-950).

